## Supplementary figures and images for "ATP increases murine neuroblastoma cell size through a PANX1- and macropinocytosis-dependent mechanism"

### Supplemental Figure 1

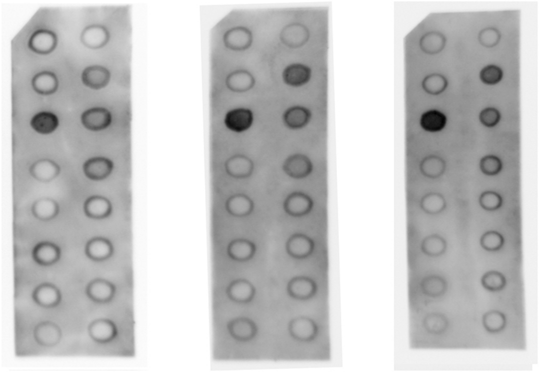

Supplemental Figure 1. Full Membrane Lipid Strip blots for Figure 3 analysis (n = 3)
